# EXD2 and WRN exonucleases are required for interstrand crosslink repair in *Drosophila*

**DOI:** 10.1101/284307

**Authors:** Pratima Chennuri, Lynne S. Cox, Robert D. C. Saunders

## Abstract

Interstrand crosslinks (ICLs) present a major threat to genome integrity, preventing both the correct transcription of active chromatin and complete replication of the genome. This is exploited in genotoxic chemotherapy where ICL induction is used to kill highly proliferative cancer cells. Repair of ICLs involves a complex interplay of numerous proteins, including those in the Fanconi anemia (FA) pathway, though alternative and parallel pathways have been postulated. Here, we investigate the role of the 3’-5’ exonuclease, EXD2, and the highly related WRN exonuclease (implicated in premature ageing human Werner syndrome), in repair of interstrand crosslinks in the fruit fly, *Drosophila melanogaster*. We find that flies mutant for EXD2 (DmEXD2) have elevated rates of genomic instability resulting from chromosome breakage and loss of the resulting acentric fragments, in contrast to WRN exonuclease (DmWRNexo) mutants where excess homologous recombination is the principal mechanism of genomic instability. Most notably, we demonstrate that proliferating larval neuroblasts mutant for either DmWRNexo or DmEXD2 are deficient in repair of DNA interstrand crosslinks caused by diepoxybutane or mitomycin C, strongly suggesting that each nuclease individually plays a role in repair of ICLs in flies. These findings have significant implications not only for understanding the complex process of ICL repair in humans, but also for enhancing cancer therapies that rely on ICL induction, with caveats for cancer therapy in Werner syndrome and Fanconi anemia patients.

## Introduction

Interstrand crosslinks (ICLs) occur between strands of the duplex DNA, preventing strand separation and thus blocking both transcription and replication of the genome. Repair of ICLs is critical to cell and organismal survival and is conducted in part by the Fanconi anemia (FA) repair pathway, which is thought to coordinate the removal and repair of covalent linkage between two DNA strands [1-3]. FA-mediated ICL repair is regulated by the S-phase checkpoint protein, ATR kinase, which phosphorylates FANCI, allowing the FA complex to monoubiquitinate FANCD2 [4,5]. Though originally thought to promote recruitment of the FANC complex to chromatin bearing a crosslink, and subsequent repair of that crosslink, monoubiquitination of FANCD2 has now been shown to occur after initial DNA recruitment [6]. Repair is thought to involve two incision events and a translesion synthesis event followed by nucleotide excision repair to remove the ICL [7]. The remaining double strand break is then repaired by homologous recombination, which requires 3’ overhang substrates and therefore is likely to involve an exonuclease acting on the opposite strand. The 5’-3’ nuclease FAN1 was identified in an RNAi screen for factors required for ICL repair following mitomycin C treatment of mammalian cells [8]. FAN1 localises to sites of crosslink damage in a manner dependent on the N-terminal UBZ domain interaction with ubiquitinated FANCD [8-10]. It has a C-terminal nuclease domain with both structure-specific endonuclease activity (that cleaves nicked and branched structures), and 5’–3’ exonuclease activity which could act in crosslink repair, although it does not appear to be involved in the trimming of DNA to allow homologous recombination [8,10-12]. The same screen suggested that EXD2, a 3’-5’ nuclease, may also be required for the response to mitomycin C, probably acting either downstream of, or in parallel to, FAN1 [8].

EXD2 has sequence similarity to the 3’-5’ exonuclease domain of human WRN (Figure 1A, also [13]), a protein with both RecQ-family helicase and DnaQ-like exonuclease domains [14-16]. Human EXD2 is predicted to exist as multiple splice variants [17] giving rise to at least two protein isoforms, with the short form predicted to lack the putative 3’-5’ exonuclease domain (Figure 1A). Here, we have chosen to investigate the role of EXD2 in *Drosophila*, in which the EXD2 orthologue is encoded by the gene *CG6744*, which produces two transcripts, CG6744-RA (1947 bp) and CG6744-RB (1850 bp) and a single predicted polypeptide of 583 amino acids. This orthologue is highly similar to human EXD2, with 40% amino acid identity and 59% similarity (Figure 1) and is referred to here as DmEXD2; it also shares significant homology with human WRN, particularly within the exonuclease domain, where key acidic residues required for nuclease activity in both human and *Drosophila* WRN are conserved in EXD2 (boxed in Figure 1A). We have previously demonstrated that DmWRNexo is indeed a *bona fide* 3’-5’ exonuclease [18], with substrate specificity highly related to that of human WRN exonuclease [18-20].

**Figure 1:**
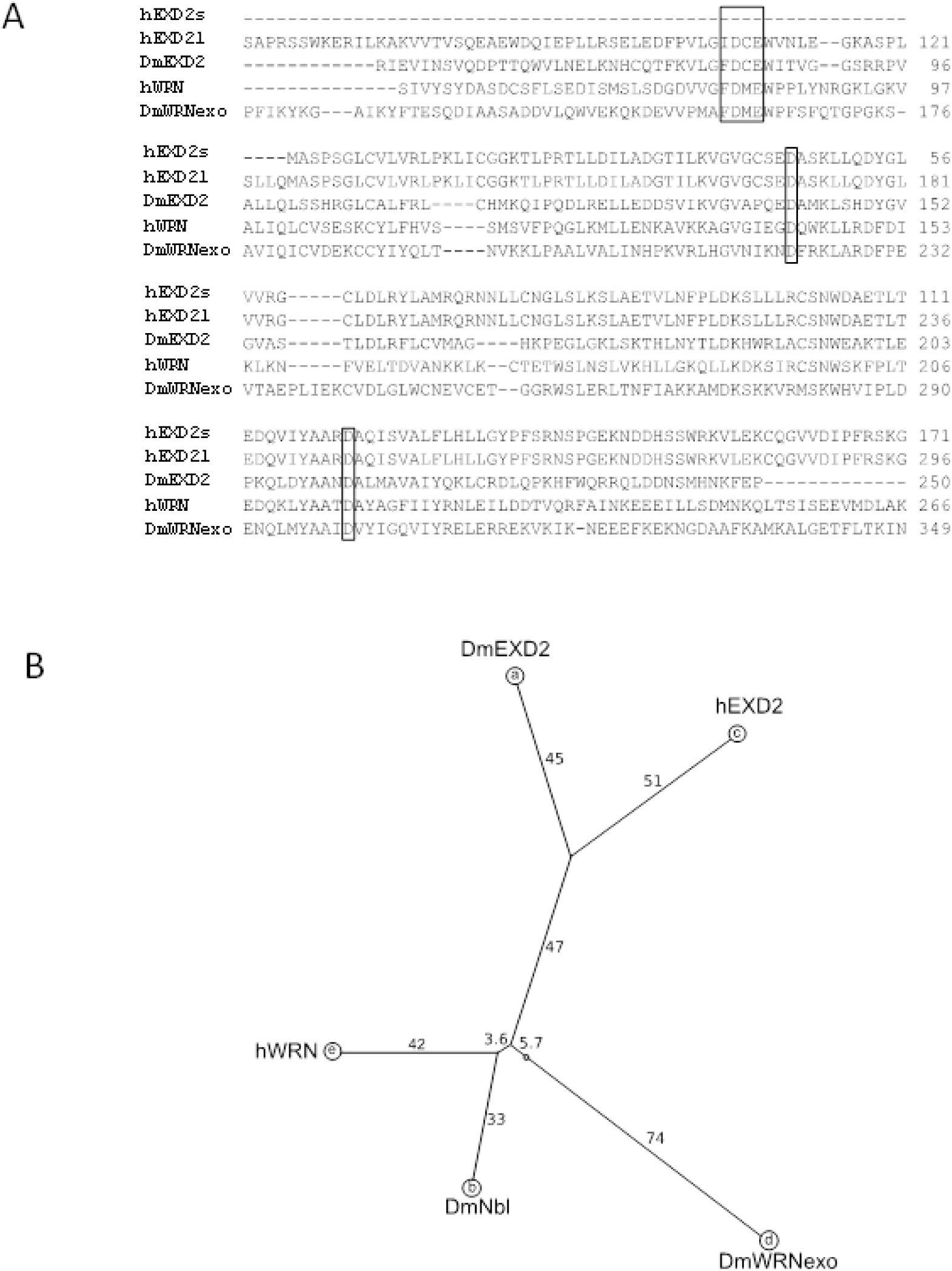
WRN and EXD2 share significant sequence similarities in the exonuclease domain. A: Alignment of the N-terminal regions of the short (497 amino acids) and long (621 amino acids) isoforms of human EXD2 (hEXD2s, hEXD2l, respectively) with Drosophila EXD2 (DmEXD2) and the related exonucleases, human WRN (hWRN) and Drosophila WRNexo (DmWRNexo). Residues critical for nuclease activity in WRN exonuclease are marked (residue numbers D82, E84, D143, D216 in human WRN). B: Unrooted phylogenetic tree to show the relationship between fly and human WRN and EXD2; note also the similarity between WRN and the fly protein Nibbler, DmNbr, a 3’-5’ exoribonuclease implicated in processing of the 3’ end of mature miRNAs (WRN has been suggested by homology to act on RNA as well as DNA [75]). The tree was generated using software developed by the Computational Biochemistry Research Group ETH Zurich.

In humans, loss of WRN function leads to the adult onset progeria Werner syndrome (WS) [21]. It is highly probable that genomic instability defects resulting from loss of WRN lead to premature cell senescence and are likely to underlie the early onset of characteristic ageing phenotypes in WS patients, including greying hair, skin changes, type II diabetes mellitus, osteoporosis, lipodystrophy and increased risk of cancer and cardiovascular disease (reviewed in [22-24]). Genomic instability arises as a consequence of WRN’s critical roles in many aspects of DNA metabolism including telomere processing [25,26] and DNA repair (e.g. [27]). Furthermore, WRN preferentially binds to DNA containing Holliday junction and/or replication fork structures [28-31], and is present at replication foci coincident with RPA and PCNA [32,33]. Cells derived from WS patients are highly sensitive to agents such as camptothecin that cause replication fork arrest or collapse, further implicating WRN in DNA replication [34-36]. Loss or mutation of WRN results in aberrant DNA replication [37-39] and hyper-recombination [40-44]. WRN associates with RAD54 family recombination proteins on treatment of cells with mitomycin C (MMC) [45], and WRN helicase inhibition potentiates toxicity in cells lacking a functional FA pathway [46]. Intriguingly, both the replication and recombination defects can be corrected by ectopic expression of the RusA Holliday junction nuclease [43,47], suggesting that it is the nuclease activity of WRN that is important in these processes within the nucleus.

Analysis of the role(s) of both EXD2 and WRN nucleases in ICL repair is important in furthering our understanding of aspects of the ICL pathway downstream of, or in parallel to, the FA complex. Here, we investigate possible roles of DmEXD2 and DmWRNexo in interstrand crosslink repair. We find that flies mutant for DmEXD2 have elevated rates of genomic instability probably resulting from chromosome breakage, in contrast to DmWRNexo mutants where excess homologous recombination is the principal mechanism of instability. Most notably, we demonstrate that flies mutant for either DmWRNexo or DmEXD2 are deficient in repair of DNA crosslinks caused by diepoxybutane (DEB) and mitomycin C (MMC), strongly suggesting that each nuclease individually plays a role in ICL repair. These findings extend our understanding of ICL repair and we propose that they highlight a novel route to enhancing ICL-dependent cancer chemotherapy.

## Materials and Methods

### Drosophila Stocks and Maintenance

DmWRNexo (*CG7670*^*e04496*^) and DmEXD2 (*CG6744*^*c05871*^) piggyBac insertional mutant strains (Bloomington *Drosophila* Stock Centre, Indiana University, http://flystocks.bio.indiana.edu) were maintained at 20°C on a standard diet of oatmeal, dextrose, cornmeal, yeast and agar supplemented with propionic acid and 10% Nipagin. Experimental crosses were maintained at 25°C.

### Wing Blade Clone Assay

Genome instability was assessed in homozygous mutant *DmEXD2*^*c05871*^ flies which were also trans-heterozygous for the cuticular markers *mwh*^*1*^ (*multiple wing hairs*, recombination map position 3-0.7) and *flr*^*3*^ (*flare*, 3-38.8). Wing blades of *mwh*^*1*^ *flr*^*+*^ *DmEXD2*^*c05871*^ */ mwh*^*+*^ *flr*^*3*^ *DmEXD2*^*c05871*^ flies were dissected from flies stored in 70% ethanol, mounted in Gary’s Magic Mountant [48] and analysed by phase contrast microscopy. Single spot (*mwh*^*1*^ or *flr*^*3*^) and twin spot clones (*mwh*^*1*^ cells adjacent to *flr*^*3*^ cells) were scored.

### Diepoxybutane Sensitivity

Balanced heterozygous virgin female and male flies mutant for *DmWRNexo*^*e04496*^ or *DmEXD2*^*c05871*^ were mated *en masse* and their progeny propagated on Instant *Drosophila* Medium (Sigma) supplemented with 0-500 μM diepoxybutane (DEB). Four independent replicate matings were conducted. Sensitivity to DEB was assessed as a deviation from the expected homozygote: heterozygote ratio in the progeny, and numbers of both male and female heterozygotes and homozygous mutants were determined. The binomial exact test for goodness-of-fit was used to ascertain statistical significance where the fly numbers were fewer than 1000, and when more than 1000, the standard chi-square test of goodness-of-fit was employed.

### Isolation and drug treatment of proliferating larval brain cells

Ventral ganglia from third instar larvae of *Oregon-R* (wild type) and larvae homozygous for *mus308*^*D2*^, *DmWRNexo*^*e04496*^ and *DmEXD2*^*c0587*^ were dissected in 0.7% saline. Tissues were transferred to HyClone SFX-Insect Culture Media (2 ventral ganglia per 100 μl) and treated with 20 μM DEB or 10 μM MMC (in culture medium) for 30 minutes during a total 4.5 hr incubation (see Figure 5A for the experimental scheme). Following treatment, tissues were collected by centrifugation, triturated into single cells and suspended in 100 μl of drug-free culture medium.

### Inverse Comet Assay for ICL Repair

Larval brain cell suspensions (see above) were mixed with 1% low melting point agarose solution at 37°C and dispersed into the well of a comet assay slide (OxiSelect™). The slides were incubated in 3 μM H_2_O_2_ for 30 minutes at 4°C to induce single strand breaks, incubated in lysis buffer (14.6% w/v NaCl, 0.1M EDTA, 1X OxiSelect™ Lysis solution (10 mM Tris, 1% Triton X-100 at pH 10)) for 45 minutes at 4°C, then transferred to alkaline solution (1.2 % w/v NaOH, 1mM EDTA) at pH>13 for 30 minutes at 4°C to denature DNA.

Electrophoresis was conducted in alkaline solution (as above) at 300 mA (1volt/cm) for 20 minutes at 4°C. Slides were washed with double distilled water for 2 minutes, 70% ethanol for 5 minutes, then stained by incubating with 1X Vista Green DNA Staining Solution (OxiSelect, Cell Biolabs, Inc.) in 1X TE buffer (10mM Tris.HCl pH 7.4, 0.1mM EDTA) for 15 minutes.

DNA migration from the nuclei was visualised as comet tails by fluorescence microscopy using the FITC filter on an Olympus BX61 motorized microscope system. A total of 10 images per treatment time point were taken and a total of 50 cells were scored per gel. Image analysis was performed with the software package CometScore (http://www.autocomet.com). All graphs were plotted by pooling data from two independent experiments per genotype, standardised against the ‘no treatment’ control (without H_2_O_2_ exposure or ICL-inducer treatment DEB or MMC, labelled H-D- and H-M- in figures). Since the data did not fit a normal distribution, non-parametric statistical tests were used to compute significance: Wilcoxon Signed Ranks test was applied to each genotype where significance of recovery at 1, 2, or 4 hours was tested against 0-hour recovery (i.e. treatment at 4h) to assess the rate and efficiency of ICL repair. The Kruskal-Wallis test was used to compute overall significance across genotypes, followed by Mann-Whitney U Wilcoxon W test for computing the significance in ICL repair of each of the test genotypes vs the wild type.

## Results

### Genome instability in DmWRNexo and DmEXD2 mutants arises through distinct mechanisms

We have previously demonstrated a requirement for DmWRNexo in maintaining genome stability in flies [49]. Given the strong similarities between DmWRNexo and DmEXD2, and the known roles of human WRN [43,50] and EXD2 [51] in homologous recombination, we first set out to establish the degree and type of genomic instability consequent upon reduction in levels of DmEXD2. We scored and analysed the appearance of recessive phenotypes in dividing wing imaginal discs in flies mutant for DmEXD2 (homozygous *DmEXD2*^*c05871*^) that are trans-heterozygous for the mutant recessive genetic markers, multiple wing hairs (*mwh*^*1*^ 3-0.3) and flare (*flr*^3^ 3-38.8).

The chromosome dynamics resulting in uncovering of mutant recessive *mwh* and *flr* phenotypes on the left arm of chromosome 3 are shown schematically in Figure 2A. Multiple wing hair (MWH) single spot clones result from either chromosome breakage proximal to *mwh* (Figure 2A (i)), or from homologous exchange in the interval between *mwh* and *flr* (Figure 2A (ii)), while flare (FLR) single spot clones result from either chromosome breakage proximal to *flr* (Figure 2A (iii)) or rare double exchange events in the interval between *mwh* and *flr* as well as proximal to *flr* (Figure 2A (iv)). In contrast, MWH-FLR twinspots (clones of MWH cells adjacent to similarly sized clones of FLR cells) necessarily arise through homologous exchange proximal to *flr* (Figure 2A (v)). An example of a twin spot is shown in Figure 2B. Wing blade cells homozygous or hemizygous for *mwh*^*1*^ bear a tuft of hairs (outlined in red, on right of in micrograph in Figure 2B) rather than a single hair in a cell-autonomous manner; flies are fully viable when homozygous. In contrast, *flr*^*3*^ causes cell-autonomous abnormal morphology of wing blade hairs that is distinct from the multiple wing hair phenotype (outlined in blue, left in micrograph in Figure 2B), and is cell-viable but organismal lethal.

**Figure 2.**
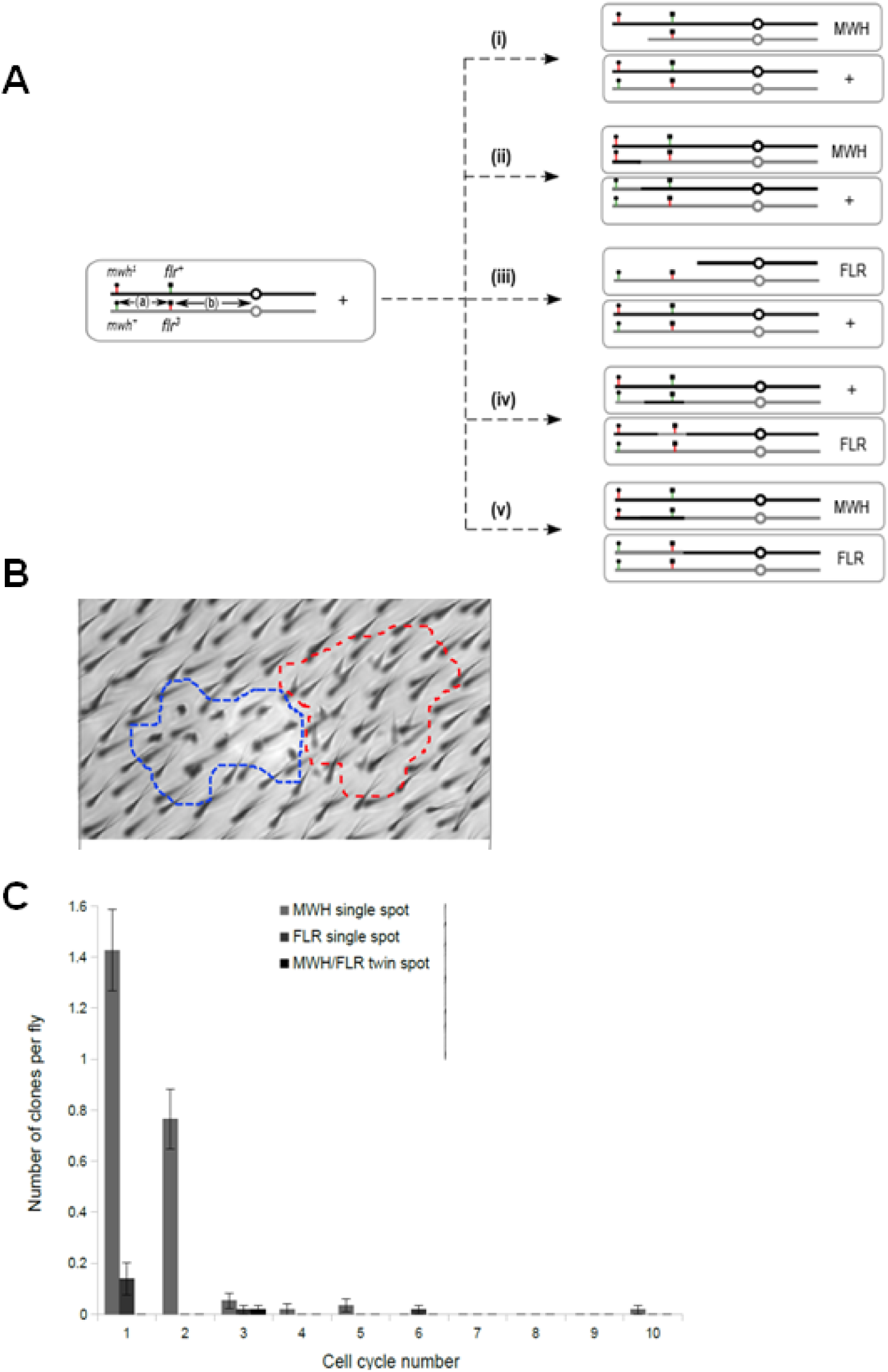
Twin spot analysis in DmEXD2 reveals chromosome breakage as major mechanism. A: Relative chromosomal positions of the markers multiple wing hairs (*mwh*) and flare (*flr*) on the left arm of Drosophila chromosome 3 are shown. Potential mechanisms for formation of wing blade clones: (i) Chromosome breakage proximal to *mwh* leads to segmentally aneuploid multiple wing hair clones, (ii) homologous exchange in the *mwh-flr* interval generates homozygous multiple wing hair clones, (iii) chromosome breakage proximal to *flr* leads to segmentally aneuploid flare clones, (iv) rare double crossover events, one between the *mwh-flr* interval and the other proximal to *flr* generate homozygous flr clones and (v) homologous exchange proximal to *flr* leads to multiple wing hair-flare twin spot clones. B. Phase contrast image of a twin spot showing *mwh* clone on the right (outlined in red) and a *flr* clone outlined in blue on the left. C: Frequencies of wing blade clones scored in *mwh*^*1*^ *DmEXD2*^*c05871*^ / *flr*^*3*^ *DmEXD2*^*c05871*^ flies, plotted against clone size in cell cycles (i.e. the number of cell divisions that have taken place since the marker was uncovered – e.g. a clone of 8 cells will reflect 3 cell cycles ie 2^3^ cells).

The number of cell divisions since the instability event occurred can be calculated from the size of the marked clone; where a breakage or recombination event is toxic such that it prevents subsequent cell division, the marker will be observed only in a single cell. The most numerous class of wing blade clones we observed were multiple wing hair single spot clones, which may arise from either chromosome breakage proximal to *mwh* or from homologous exchange between *mwh* and *flr* (Figure 2A (i) or (ii)). Multiple wing hair-flare (MWH-FLR) twin spot clones can only arise from homologous exchange and are much rarer. We observed a high frequency of MWH clones in DmEXD2 mutant flies indicating genomic instability, compared with MWH-FLR twinspots, which are very rare in these mutants (Figure 2C), strongly suggesting that the principal mechanism of genomic instability is chromosome breakage with subsequent loss of the acentric fragments. This contrasts with our finding in flies mutant for DmWRNexo, where the principal mechanism of genomic instability is excessive homologous recombination [49], and strongly suggests that the two related nucleases act in different molecular pathways.

### DmWRNexo and DmEXD2 mutant flies are sensitive to the ICL inducer diepoxybutane

Given the difference in the mechanism of genomic instability consequent on WRNexo or EXD2 loss, we wished to further explore their roles in maintaining genomic integrity. We have previously demonstrated that flies hypomorphic for WRNexo are hypersensitive to the topoisomerase I inhibitor camptothecin [49], consistent with a requirement for WRN exonuclease during normal or perturbed DNA replication [38,39,47]. Human WRN helicase has also been suggested to act in DNA interstrand crosslink repair [52], supported by studies involving inhibition of WRN helicase [46]. To date, however, no study has directly assessed a possible role of WRN exonuclease in ICL repair because of the difficulty in isolating the nuclease from the helicase activity - dominant negative mutation in one activity is likely to impact on the other. Since flies encode the WRN exonuclease on a genetic locus discrete from the putative WRN helicase [49], we were able to assess the role of WRN exonuclease in ICL repair, and compare it with the related nuclease EXD2.

We tested the sensitivity of flies mutant for either DmWRNexo or DmEXD2 to the crosslinking agent diepoxybutane (DEB), administered in the diet at concentrations much lower than previously shown to highlight defects in DNA repair pathways [53]. The ratio of homozygotes to heterozygotes in the progeny was assessed for each DEB concentration (0-0.5 mM): any deviation from a homozygote:heterozygote ratio of 1:2 in a monohybrid balanced heterozygote cross (the balancer chromosomes carry recessive lethal alleles) indicates sensitivity of the homozygous mutant flies to DEB treatment.

Our data (Figure 3 and Supplementary Table S1) demonstrate that flies homozygous for either *DmWRNexo*^*e04496*^ or *DmEXD2*^*c05871*^ are extremely sensitive to DEB; *DmWRNexo*^*e04496*^ mutants show slightly greater sensitivity than *DmEXD2*^*c05871*^ mutants. No *DmWRNexo*^*e04496*^ homozygotes survived at a DEB concentration of 0.2 mM or higher; note that this is ten-fold lower than the dose to which *mus201* and *mei-41* mutant flies are reported to be sensitive [53]. Supplementary table S1 catalogues the number of flies eclosing for each genotype at each concentration of DEB, together with the probability that the ratio of homozygotes to heterozygotes differs significantly from the predicted value of 1:2 (i.e. ratio of 0.5 in Figure 3 and Table S1). These results confirm that both DmWRNexo and DmEXD2 are required for resistance to DNA crosslinking agents such as DEB, suggesting a role in ICL repair. We find no sex differences in sensitivity of either DmWRNexo or DmEXD2 mutants to DEB (Figure 3B and 3C respectively). The hypersensitivity of *DmWRNexo*^*e04496*^ mutants (strongly hypomorphic for the WRN exonuclease) to a crosslink inducer is an intriguing finding as it suggests an important role for WRN’s exonuclease activity in ICL repair, in addition to the previously suggested ICL repair role of the helicase domain [46,52]. The DEB sensitivity of DmEXD2 mutants is particularly intriguing, as it provides the first *in vivo* confirmation of previous *in vitro* suggestions that implicate EXD2 in ICL repair [8].

**Figure 3.**
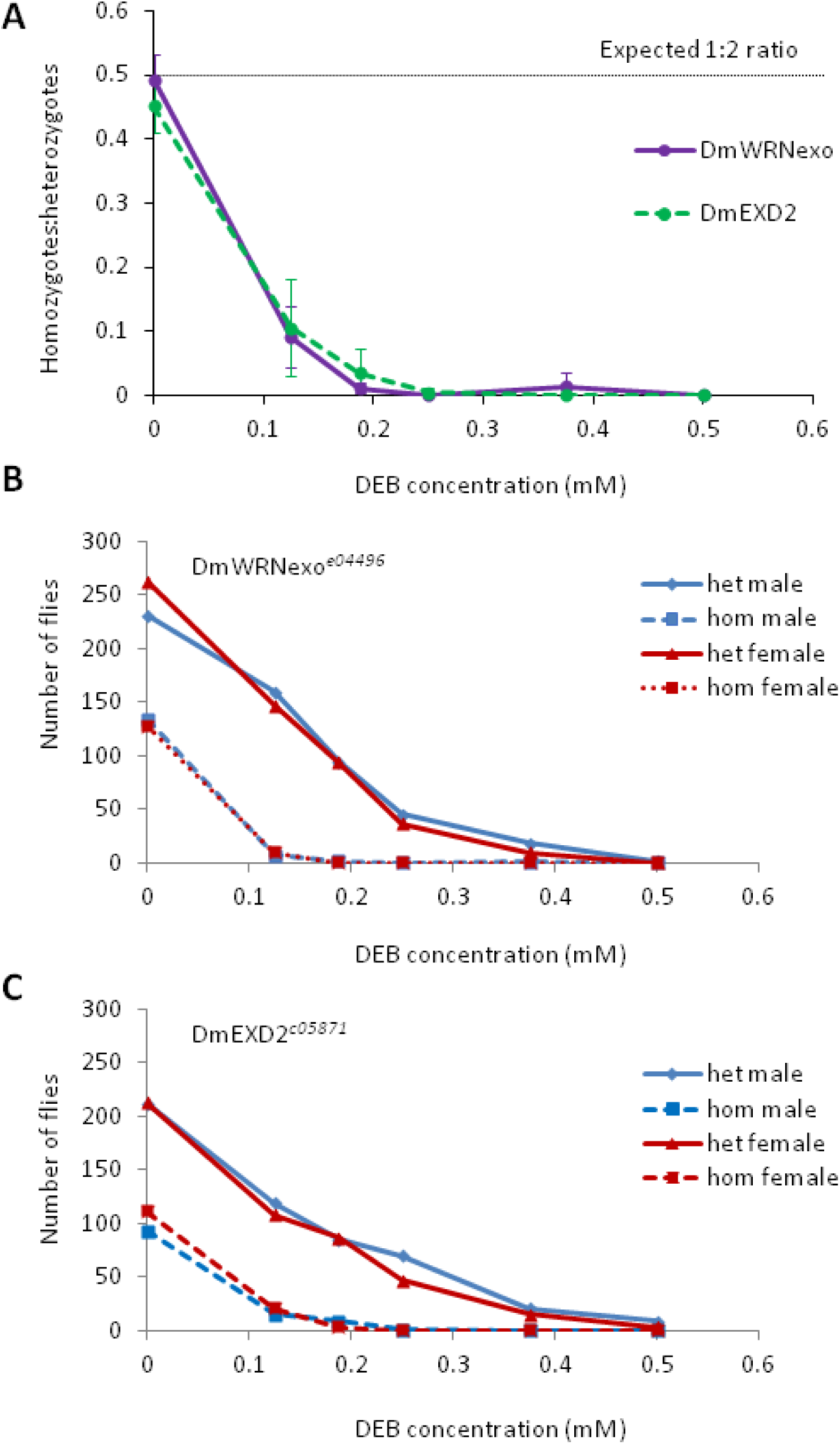
DmEXD2 and DmWRNexo mutants are very sensitive to the ICL inducer diepoxybutane. A: Balanced heterozygous flies with *DmEXD2*^*c05871*^ or *DmWRNexo*^*e04496*^ were mated and progeny raised on medium supplemented with diepoxybutane (DEB) at the indicated concentrations. The ratio of homozygotes to heterozygotes (ie *DmEXD2*^*c05871*^/*DmEXD2*^*c05871*^ *to DmEXD2*^*c05871*^/*+* or *DmWRNexo*^*e04496*^/ *DmWRNexo*^*e04496*^ to *DmWRNexo*^*e04496*^/*+*) was calculated and plotted against DEB dose, as an indicator of relative sensitivity (note that due to the lethal balancer, flies that are homozygous wild-type for DmWRexo or DmEXD2 would not be observed in these experiments). The threshold line at 0.5 represents the expected homozygote to heterozygote ratio for control treatments (i.e. no DEB in medium). Significant decreases in this ratio imply significant sensitivity of homozygous mutant flies to DEB-induction of ICLs (see Supplemental Table S1 for statistical analysis). Data shown are means of 4 independent experiments; error bars = +/- standard deviation B, C: Pooled data from 2 independent experiments showing number male (blue) and female (red) mutant flies following DEB treatment for strain DmWRNexo^*e04496*^ (B) and *DmEXD2*^*c05871*^ (C). Heterozygotes are represented by continuous lines, and homozygotes by dotted lines.

### Defective ICL repair contributes to DEB sensitivity in DmWRNexo and DmEXD2 mutant flies

To confirm that DEB sensitivity was due to defects in ICL repair, persistence of ICLs following recovery from acute DEB exposure was assessed by an inverse comet assay, which measures the ability of cells to recover from drug-induced ICL damage. In this assay, ICLs were induced by treatment with DEB, then following various recovery periods, nuclei were embedded in low melting point agarose and exposed to H_2_O_2_ to generate nicks in the DNA. Intact DNA is retained in the nucleus on electrophoresis (Figure 4A (i) undamaged) while DNA that is subsequently nicked by the peroxide treatment will migrate from the nuclei on electrophoresis (Figure 4A, (ii) nicked DNA). Treatment with agents that induce ICLs prior to H_2_O_2_ treatment results in retention of DNA within the nucleus as crosslinked DNA is unable to migrate out of the nucleus (Figure 4A, (iii) ICL formation), while repair of the ICLs will allow nicked DNA to migrate (Figure 4A (iv) ICL repair). Hence a larger comet tail moment (after H_2_O_2_ nicking) is indicative of greater ICL repair. The presence of a genetic mutation that impedes ICL repair will result in shorter comet tails in comparison with wild type, signifying the importance of that gene product in the repair of ICLs. In order to determine optimal concentrations of H_2_O_2_, DEB and MMC, standard comet assays were also performed, verifying that peroxide treatment results in greatly extended comet tails, with the bulk of the nuclear DNA extruding from the cell on electrophoresis (Figure 4B). By contrast, DEB treatment shortens the comet tail moment in a dose-dependent manner (Figure 4C). On the basis of these and related optimisation studies (supplementary Figure S1 and data not shown), concentrations of 3μM H_2_O_2_, 20 μM DEB and 10 μM MMC were chosen for subsequent experiments.

**Figure 4.**
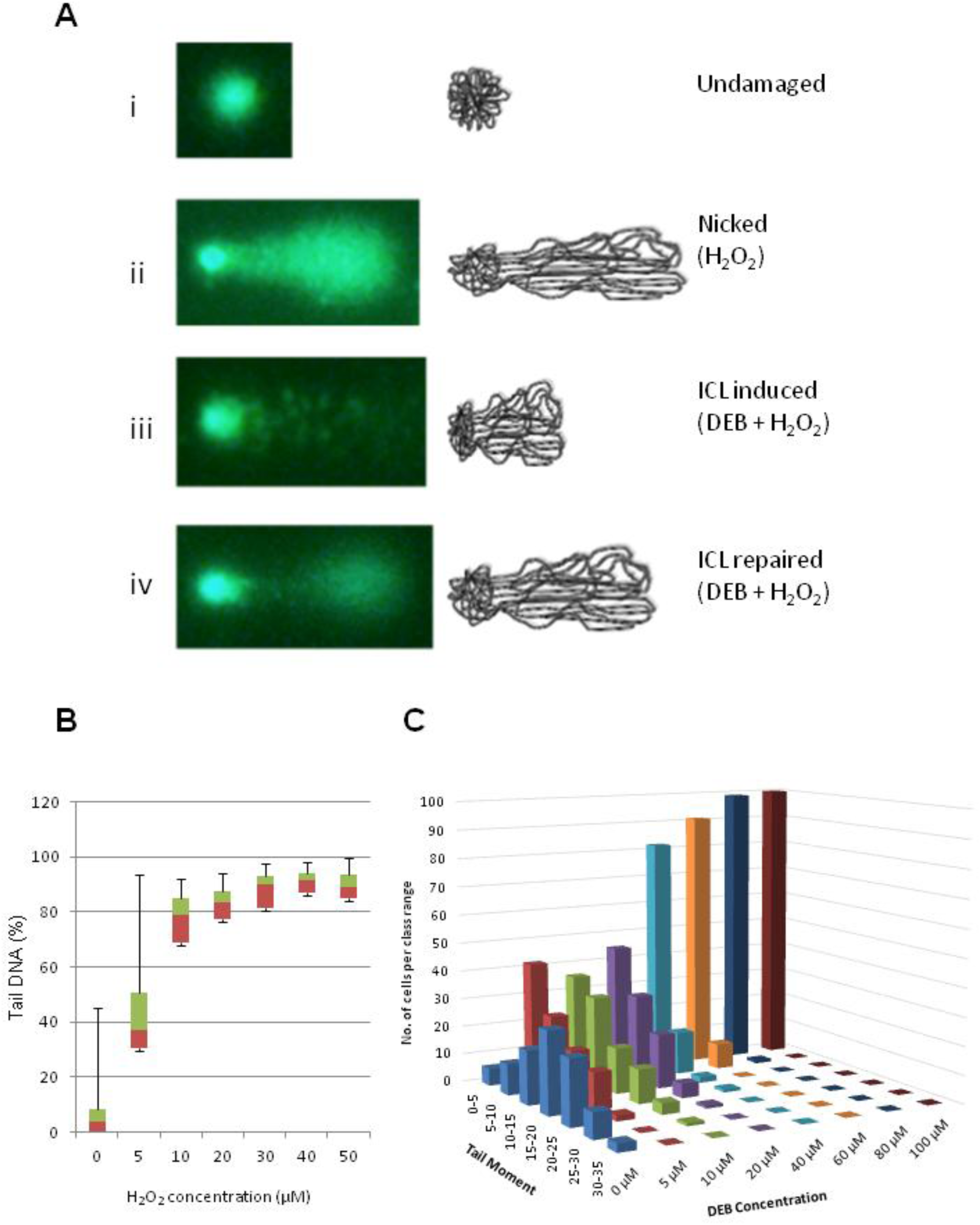
Inverse comet assay for induction of and recovery from ICL damage. A: Schematic to show outcomes of the inverse comet assay, whereby a comet tail is only elicited if DNA is free of interstrand crosslinks and is nicked by peroxide treatment. (i) undamaged nuclei with intact DNA retain the bulk of the DNA within the nucleus upon electrophoresis. (ii) Treatment with H_2_O_2_ induces DNA nicking and breakage such that DNA extrudes from the nucleus on electrophoresis to form an extensive comet tail. (iii) Treatment with an ICL inducer (eg DEB) prior to H_2_O_2_ treatment reduces the length of the comet tail as cross-linked DNA cannot migrate from the nucleus. (iv) repair of ICLs prior to H_2_O_2_ treatment leads to increased tail length compared with immediately after ICL induction. B: Determination of peroxide concentrations suitable to elicit a measurable comet tail. Box plots showing 25^th^, median, 75^th^ percentile values of % DNA present in the comet tail for DmEXD2^*c05871*^ larval neuroblast nuclei treated with varying concentrations of hydrogen peroxide; whiskers represent 0 and 100%. Standard comet conditions were used. C: Determination of DEB concentrations necessary to retain DNA in nuclei under inverse comet conditions without recovery period. The frequency plot shows combined data from two independent biological repeats for DmEXD2^*c05871*^ larval neuroblast nuclei. 50 nuclei were counted for each experiment for each drug concentration giving a combined total of 100 nuclei per DEB concentration.

We then exposed proliferating Drosophila cells isolated from neural tissue of larvae of various genotypes (wild type *Oregon-R* and homozygotes mutant for either *DmWRNexo, DmEXD2* or *mus308*) to the ICL inducer DEB, with variable recovery times prior to inverse comet assay, according to the schedule depicted in Figure 5A. The impact of treatment is shown for each mutant genotype individually compared with *Oregon-R* wild type in Figure 5B-D; Figure 5E shows all genotypes tested in these experiments in the same graph for ease of comparison. Controls treated with peroxide but not DEB are designated H+D-; controls without either treatment are indicated by H-D-.

Neuroblasts from wild type larvae (*Oregon R*) showed significant increase (p<0.001) in the measured comet tail moment when allowed to recover from DEB treatment for >1 hour, indicating rapid and robust levels of ICL repair (Figure 5B-E). After 4 hours of recovery, no further increase in tail moment was detected, representing ∼60% of that seen for controls treated only with peroxide and not DEB (H+D- in Figure 5B-E). Given the toxicity of DEB even at the low concentrations used here (see also Figure 3A and 3B), this failure to achieve levels seen in controls may reflect death of some of the DEB-treated cells during the course of the experiment. That the comet tail results from experimentally-induced nicking by H_2_O_2_ can be clearly demonstrated by the lack of DNA migration from control nuclei not exposed to peroxide (Figure 5, H-D-). A further control with peroxide but without DEB treatment (Figure 5B-E, H+D) demonstrates that nicking and comet tail formation are similar irrespective of genotype, providing verification of the experimental strategy. Notably, we find that larval neuroblasts from hypomorphic DmWRNexo mutants (*i.e. DmWRNexo*^*e04496*^) showed defects in ICL repair (Figure 5B); the lack of repair was highly significant and equivalent to that seen in the positive control DNA polymerase theta mutant, *mus308*^*D2*^ (Figure 5D), which has been reported to be required in ICL repair [54]. We observed no significant increase in comet tail moment over the recovery time periods following DEB treatment, confirming the absence of ICL repair in the *mus308*^*D2*^ mutant neuroblasts (Figure 5D). Moreover, larval neuroblasts derived from *DmEXD2*^*c05871*^ homozygotes also showed a defect in ICL repair; though a slight increase in comet tail moment was detected at the 1-hour recovery time point, no further increase was observed with longer recovery periods, even after 4 hours following removal of DEB (Figure 5C), strongly suggesting that these cells cannot efficiently eliminate ICLs. Pairwise comparisons of ICL repair efficiency of each of the test genotypes vs wild type (*Oregon-R*) revealed a severe defect in ICL repair on mutation of MUS308, DmWRNexo or DmEXD2 proteins (Supplementary Table S2). The Wilcoxon Signed Ranks test was used to test significance of recovery patterns over time relative to the 0-hour time point for each strain (Supplementary Figure S2). Together these data show that both DmWRNexo and DmEXD2 are required for efficient ICL repair, at least in the highly proliferating cells in larval fly brains.

### Both DmWRNexo and DmEXD2 are required for efficient repair of ICLs induced by mitomycin C

Since crosslink-induced structural distortion influences the recruitment of specific protein components from different DNA repair pathways (reviewed in [55]), the inverse comet assay was repeated with the ICL inducer mitomycin C (MMC), which produces structural distortions that are different from those produced by DEB. An experimental approach identical to that of DEB was adopted, exposing larval neuroblasts to MMC for 30 minutes with variable recovery periods (see schematic Figure 5A). As with cells treated with DEB, MMC treatment led to decreased comet tail moments as a consequence of formation of interstrand crosslinks. In the wild type *Oregon R* neuroblasts, significant and sustained increase in mean comet tail moment was observed with increasing recovery time following 30-minute exposure to MMC (Figure 6, all panels), indicating repair of the cross-links. In contrast, neuroblasts obtained from larvae mutant for either DmWRNexo (Figure 6A) or DmEXD2 (Figure 6B) showed impaired recovery from MMC-induced ICL damage, to a similar degree as neuroblasts mutant for *mus308*^*D2*^ (Figure 6C), known to be defective in ICL repair. Pair-wise comparisons of mean comet tails moment show highly significant differences between the wild type *Oregon-R* strain and all mutant strains tested (Supplementary Table S3). Taken together, our data demonstrate that ICL repair is defective in fly larvae mutant for either DmWRNexo or DmEXD2, irrespective of the ICL-inducing agent used.

## Discussion

Investigations into interstrand crosslink repair generally focus on the complex and still poorly understood Fanconi anemia pathway. However, steps parallel to, or downstream of, FA complex intervention are also critically important not only in understanding how ICL repair is mediated in cells, but also in informing on design of pharmacological modulation, either to promote or, perhaps more excitingly, to inhibit the repair of ICLs in synthetic lethality treatments for cancer using ICL-inducing therapies. Such synthetic lethality therapies, first suggested in 1997 [56], have shown great promise in breast cancer where BRCA-mutant cells treated with PARP inhibitors are exquisitely sensitive to agents inducing double strand breaks (reviewed in [57]).

**Figure 5:**
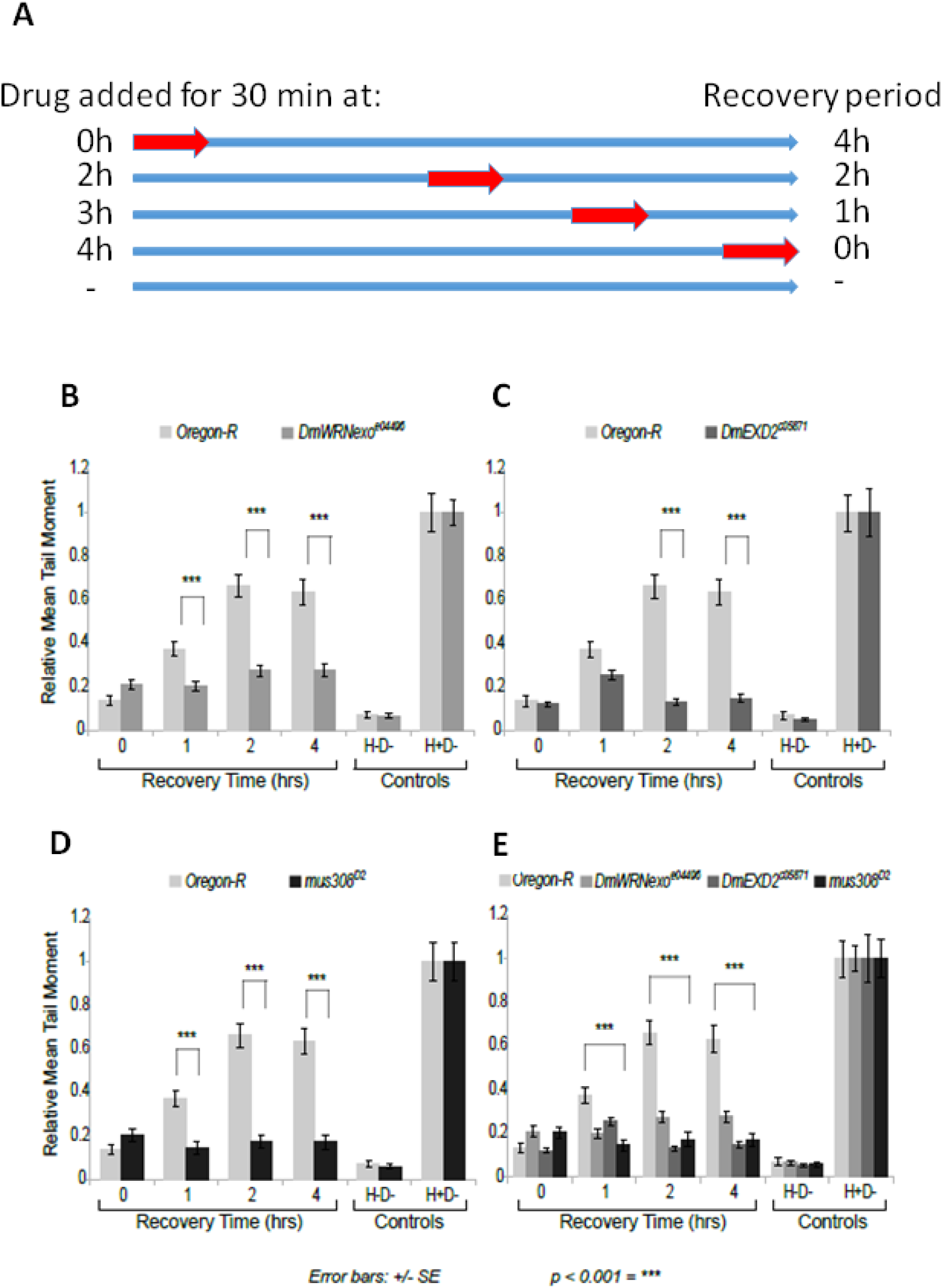
Cells mutant for DmWRNexo or DmEXD2 show impaired repair of DEB-induced ICLs. A: Schematic showing experimental regime for 30-minute treatment with 20 μM diepoxybutane DEB (red arrows) and recovery period over a total experimental time of 4.5 hours. B: ICL repair capacities of *DmWRNexo*^*e04496*^ lies versus wild type (*Oregon-R*) flies; C: comparison of *DmEXD2*^*c0587*^ versus wild type; D: known ICL repair-deficient (*mus308*^*D2*^) mutants compared with wild-type; E: all genotypes shown together for ease of comparison. Controls include H-D- i.e. not treated with either H_2_O_2_ or DEB, and H+D- i.e. treated with peroxide but not DEB. An increase in comet tail moment following recovery periods after DEB treatment indicates efficient ICL repair. The data within each genotype were standardized to the H+D- control and statistical significant differences between mutant strains and wild type Oregon-R was calculated using the Mann-Whitney U Wilcoxon W test. *** represents p<0.001 (see Supplementary Table S2 for more details). Error bars represent standard error.

**Figure 6.**
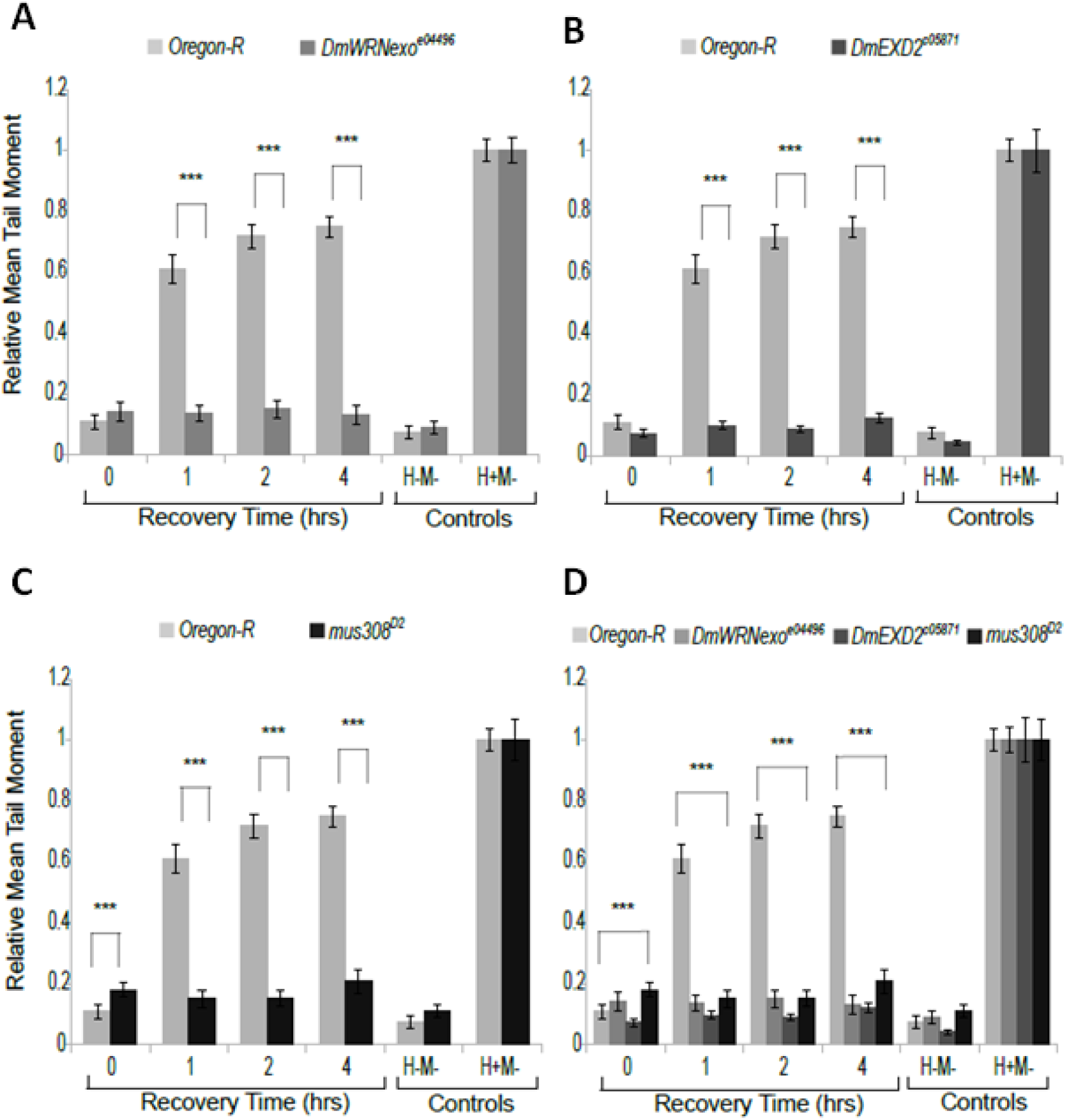
DmWRNexo and DmEXD2 are required for repair of MMC-induced ICLs. An inverse comet assay was conducted on larval neuroblasts following treatment with ICL-inducer 10 μM mitomycin C for 30 minutes followed by variable recovery periods (see schematic in Figure 5A). A: ICL repair capacities of wild type (*Oregon-R*), compared with *DmWRNexo*^*e04496*^ mutant flies. B: comparison of homozygous mutant *DmEXD2*^*c0587*^ larval neuroblasts vs wild type. C: Known ICL repair-deficient (*mus308*^*D2*^) larval neuroblasts compared with wild type. D: All genotypes together. Controls are without either peroxide or MMC treatment (H-M-) and with peroxide but without MMC (H+M-). Data within each genotype were standardized to the H+M- control and significance was calculated as for Figure 5. *** represents p<0.001 (See Supplementary Table S3 for more detail). Error bars represent standard error.

The progeroid WRN helicase/exonuclease has been implicated in ICL repair following observations of sensitivity of patient-derived cells to ICL-inducing agents (e.g. [58]). Additional nucleases, particularly FAN1 and EXD2 have been proposed to act during ICL repair, but such conclusions are based on *in vitro* data using either purified proteins and DNA substrates [55] or RNAi screens in cultured cells [8]. Here, we have examined ICL repair in a whole organism, demonstrating requirement for both WRN and EXD2 nucleases in the ICL repair pathway. Drosophila provides significant advantages over cell culture models: firstly, genetic markers allow us to determine not only the degree of genomic instability resulting from loss of function of these nucleases, but also the type of chromosomal damage caused. Secondly, invertebrates encode the functionalities of WRN helicase and exonuclease on separate genetic loci [59] allowing a genetic dissection of their respective roles without the hindrance of dominant negative mutation effects seen in human WS cells.

Our results presented here using genetic markers as readouts of chromosomal instability phenotypes not only confirm our earlier findings of spontaneous genomic instability on hypomorphic mutation of either DmWRNexo [49] or DmEXD2 [13] but allow us to conclude that loss of WRN exonuclease results in excess homologous exchange while decreased DmEXD2 activity instead results in chromosomal breakage. Chromosome breakage leads to growth impairment due to segmental aneuploidy and consequently fewer FLR single spot clones would be expected than MWH single spot clones in the wing blade assay shown in Figure 2, as terminal deletions uncovering *flr* would on average be larger than those uncovering *mwh*; by contrast, a high frequency of MWH-FLR twinspots should be observed if the mechanism of instability is excessive homologous recombination. Given that DmWRNexo mutants show a high frequency of wing blade clones and that several of these have arisen from multiple cell divisions since the original mutation, we can with confidence conclude that the cells are not suffering from aneuploidy and instead that the phenotype must have arisen from reciprocal exchange, while the lower frequency of clones seen in DmEXD2 mutants may reflect a role for DmEXD2 in other types of repair that occur outside S phase (i.e. in G1 or G2 phases). This difference in both frequency and mechanism of genomic instability between flies mutant for DmWRNexo [49] and DmEXD2 (data presented in this paper), suggests that the two highly related enzymes function in distinct molecular pathways and possibly in temporally distinct phases of the cell cycle, though we cannot yet rule out the possibility of partial redundancy.

To date, studies on WRN’s involvement in ICL repair have focussed on its helicase activity: FANCD2 knockout cells become more sensitive to MMC when treated with a small molecule inhibitor of WRN helicase (NSC 617145), while exposure to NSC 617145 and MMC led to elevated levels of Rad51 foci [46] suggesting that WRN helicase is required for later steps of HR at ICL-induced DSBs [45]. However, it has been so far unclear if WRN exonuclease plays any part in ICL repair. We addressed this using the Drosophila *DmWRNexo*^*e04496*^ mutant, as this is severely hypomorphic for DmWRNexo [49], but has intact helicase activity (we find synthetic lethality when both the *DmWRNexo* and the putative helicase encoded by *mus309* are mutated – data not shown). We also utilised flies hypomorphic for DmEXD2, which is highly related to WRN, to assess EXD2 involvement in ICL repair.

It has been proposed that the Fanconi anemia (FA) pathway is the preferred pathway of ICL repair in cells undergoing active DNA replication (i.e. in S phase), which favours homologous recombination-mediated repair of ICL-induced DSBs in collapsed replication forks [60]. Notably, we have shown that replication forks collapse without WRN function [38] and that restoration of a Holliday-junction nuclease activity can overcome this defect [47]. Hence it is likely that WRN exonuclease may play a role in ICL repair, possibly during the HR step, particularly in highly proliferating cells. We therefore chose to study larval neuroblasts as these represent a proliferative population that is accessible to experimental intervention following dissection of larval neural tissues. We challenged proliferating Drosophila cells derived from both DmWRNexo and DmEXD2 mutant strains with two ICL inducers, diepoxybutane and MMC, since these agents cause different degrees of helical distortion in duplex DNA and may therefore trigger divergent ICL repair pathways. We clearly demonstrate that both EXD2 and WRN exonuclease are required for repair of ICLs (irrespective of mode of induction), since cells lacking either activity fail to remove cross-links, as demonstrated in our inverse comet assay.

We find notable differences in the recovery of larval neuroblasts on exposure to DEB compared with MMC. In particular, DmEXD2 mutants showed low but significant levels of repair following a one hour recovery period after exposure to DEB (Figure 5C) but no recovery when exposed to MMC (Figure 6B). In the absence of a recovery period, the mutant neurobasts treated with DEB were much worse in terms of repair than untreated controls, while the ICL repair-deficient *mus308*^*D2*^ homozygous neuroblasts were instead slightly more sensitive to DEB than to MMC. These differences might be explained (at least in part) by the chemical, structural, and mechanistic differences in the induction of DNA damage by the individual molecules, although the exact mechanism remains to be determined. Both DEB and MMC are bifunctional alkylating agents and form monoadducts and/or abasic sites in addition to ICLs (biadducts), although their cytotoxicity is attributed to ICLs [61-63]. These perturbations typically result in single-stranded DNA breaks, the rate of formation of which might differ between the two alkylating agents, resulting in the observed differences. Specifically, ICLs produced by DEB result in substantial DNA helix distortion to the magnitude of ∼34° per lesion towards the major groove [64] while those produced by MMC are minimal [65]. The extent of helical distortion has been shown to influence the efficiency of nuclease activity near ICLs, with the less-distorting lesions showing a ∼3-fold lower incision frequency [66]. This might effectively make MMC-induced ICLs harder to nick, resulting in a more efficient readout of ICL repair deficiency in the mutant genotypes vs the wild type (Figure 6). Despite these expected drug-dependent differences, the finding that ICL repair is defective in both *DmWRNexo*^*e04496*^ and *DmEXD2*^*c05871*^ single mutants is highly suggestive that each nuclease is necessary in order to carry out efficient ICL repair. We have therefore, for the first time, demonstrated that both EXD2 and WRN exonucleases are required for efficient repair of ICLs.

Crosslinking agents used in cancer therapy can also drive mutagenesis and yield secondary malignancies. Since crosslinker-induced cytotoxicity and mutagenesis can be altered differently by different DNA repair pathways, it is important to understand all possible mechanisms of ICL toxicity. An important consideration in therapeutic regimens involving combination therapy is the spectrum of DNA damage induced by specific crosslinking agents, where agents with complementary spectra of DNA damage can derive maximum benefit [67]. ICL-inducing agents may be particularly effective cancer treatment in the presence of FA gene mutations, so in tumours with intact ICL repair genes, an alternative treatment strategy would include a combination of ICL-inducing agents together with inhibitors of ICL repair pathways [68]. Inclusion of inhibitors of WRN and/or EXD2 as possible therapeutic agents to increase the efficacy whilst reducing the required dose of crosslinking agents such as cisplatin might have great potential for developing safer and more efficient treatment strategies for cancer, as ICL-inducers at currently used doses are highly toxic to normal as well as cancer cells leading to serious side-effects [46].

Our findings also have implications for human ageing processes. Chronic accumulation of DNA damage, both from prolonged exposure to low-level environmental genotoxins and endogenous damage, and from errors in replication, repair and recombination, can trigger a p53-dependent DNA damage response that ultimately result in cellular senescence. Werner patient cells show both increased genomic instability (especially at the telomeres) and accelerated onset of cellular senescence, suggesting that their inability to effectively tackle DNA damage may drive senescence. While the rate of spontaneous ICL formation is currently unknown, the evolution of pathways for efficient repair suggests that it is not trivial. We suggest that better understanding of ICL repair pathways involving WRN or EXD2 may lead to the discovery of targets and small molecules that may have the potential to alleviate or delay the adverse effects of cell senescence caused by some genotoxic stresses – as accumulation of senescent cells is detrimental to tissue integrity and function [69-73], such agents would have potential to alleviate diseases of ageing.

We conclude that both WRN and EXD2 could serve as important targets for treating age-associated adverse effects including some types of cancers. Combination synthetic lethality therapy should aid development of more efficient treatment strategies with relatively milder side effects due to reduction in doses of harmful non-specific genotoxic chemotherapeutic agents. However, our findings also bring an important caveat for patients with FA or progeroid Werner syndrome, (and their attending physicians). It is already known that in FA patients, lack of repair of therapy-induced ICLs in non-cancerous cells can lead to dangerous consequences including hematopoietic apoptosis, dysplasia of remaining cells and subsequent cancer, as well as increased cancer incidence arising from opportunistic viruses, since ICL-inducers lead to such patients becoming immune-compromised [74]. Moreover, Werner syndrome patients are likely to be hypersensitive to ICL-inducing agents, so it is vital to caution against use of such drugs without much more thorough understanding of likely systemic effects. We therefore propose that ICL-inducers together with transient blockade of ICL-repair pathways through pharmacological WRN or EXD2 inhibition may be beneficial for cancer patients with otherwise normal DNA repair pathways, but that ICL-inducing therapies should be avoided in individuals already compromised for DNA repair.

## Acknowledgements

We are grateful to the BBSRC for funding that led to this project including grant numbers [BB/E000924/1] to LSC and [BB/E002072/1] to RDCS, the Glenn Foundation for Medical Research for a Glenn award to LSC, and the Open University for a studentship award to RDCS that supported PC.

---

**Table S1.**
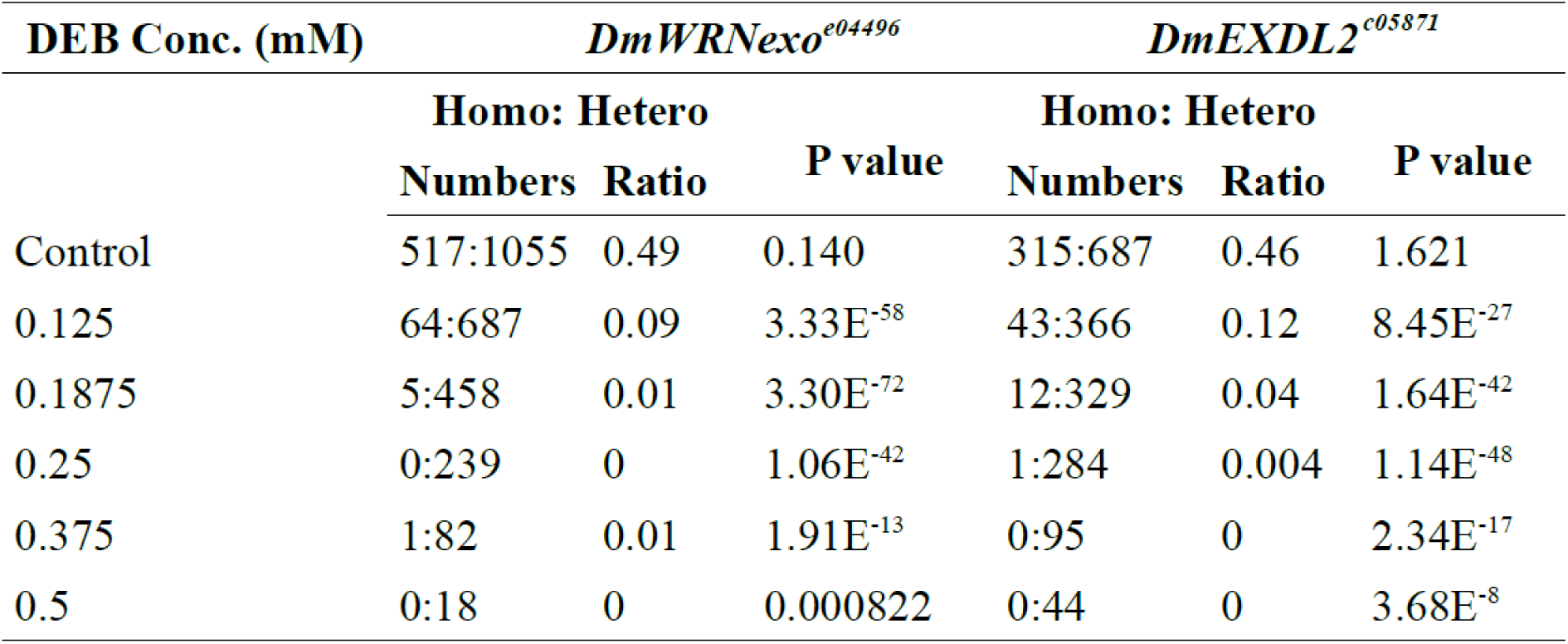

**Supplementary Figure S1.**
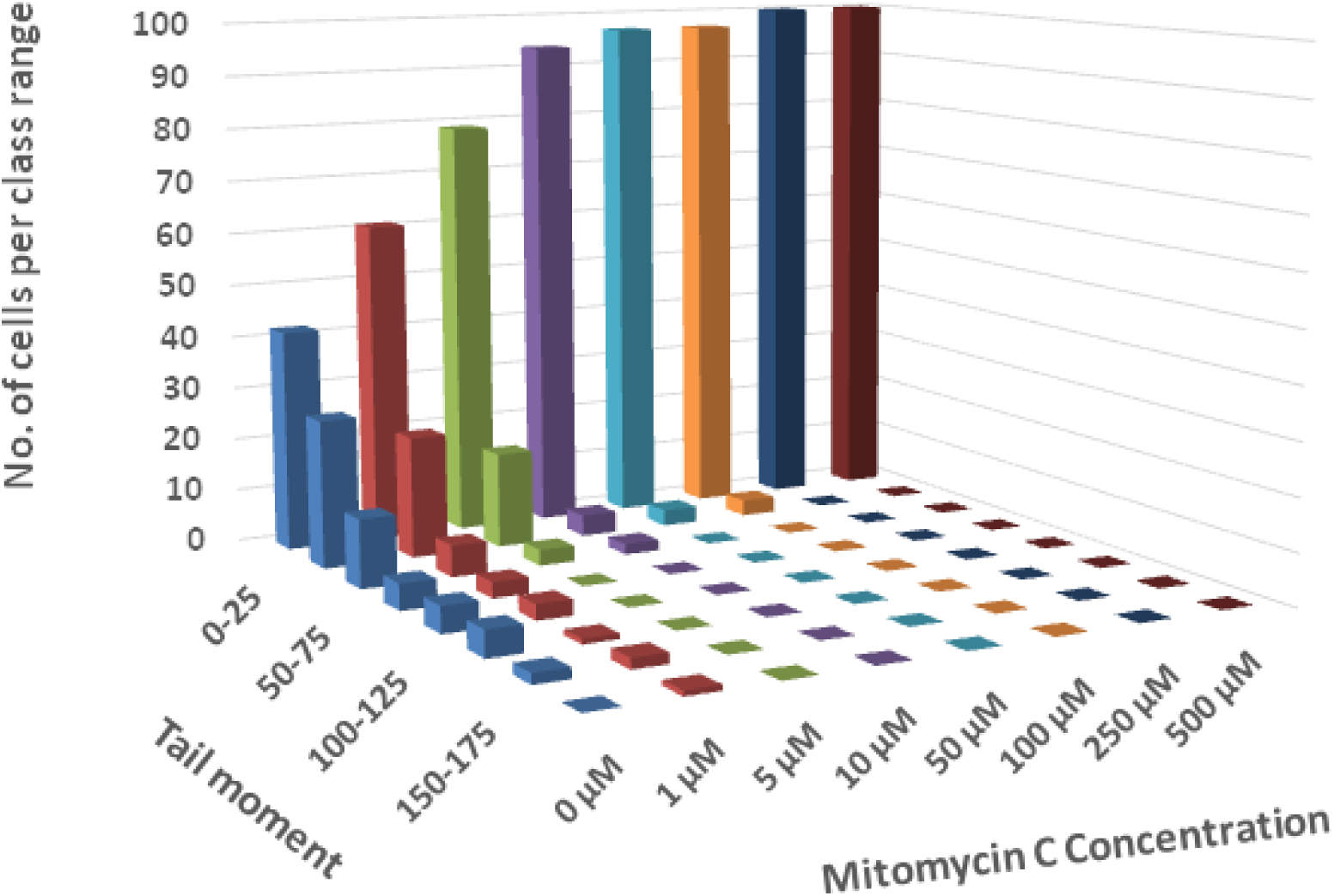
Determination of optimal mitomycin C (MMC) concentration to generate interstand DNA cross-links. Wild type Oregon-R larval neuroblasts were exposed to MMC at the shown concentrations for 30 minutes and a standard comet assay conducted (ie without hydrogen peroxide nicking). Data shown are pooled from two independent experiments where 50 nuclei were measured for each drug condition, giving a total 100 nuclei for each dose.

**Supplementary Table 2.** Statistical significance of repair efficiency of DEB induced ICLs. Analysis of difference in comet tail moment at various recovery times following DEB-induced ICLs in homozygous mutant fly larval neuroblasts versus using Oregon-R wild type controls. The Mann-Whitney U Wilcoxon W Test was used for pairwise comparisons between each mutant genotype and the wild type, and overall significance tested using the Kruskal Wallis Test package, using SPSS. H-D- = control without either hydrogen peroxide (nicking) or DEB (cross-linking) treatment. H+D- = control with peroxide but without DEB. *** in Figure 5 indicates p<0.001.

|  | *Oregon-R (Pairwise and overall significance)* |  |  |  |  |  |
| --- | --- | --- | --- | --- | --- | --- |
|  | 0 hr | 1 hr | 2 hr | 4 hr | H-D- | H+D- |
| *DmWRNexo*<sup>e04496</sup> | 0.001 | 0.000 | 0.000 | 0.000 | 0.243 | 0.306 |
| *DmEXD2*<sup>c05871</sup> | 0.017 | 0.124 | 0.000 | 0.000 | 0.104 | 0.216 |
| *mus308*<sup>D2</sup> | 0.001 | 0.000 | 0.000 | 0.000 | 0.017 | 0.604 |
| Overall | 0.001 | 0.000 | 0.000 | 0.000 | 0.160 | 0.225 |

**Supplementary Table S3.**
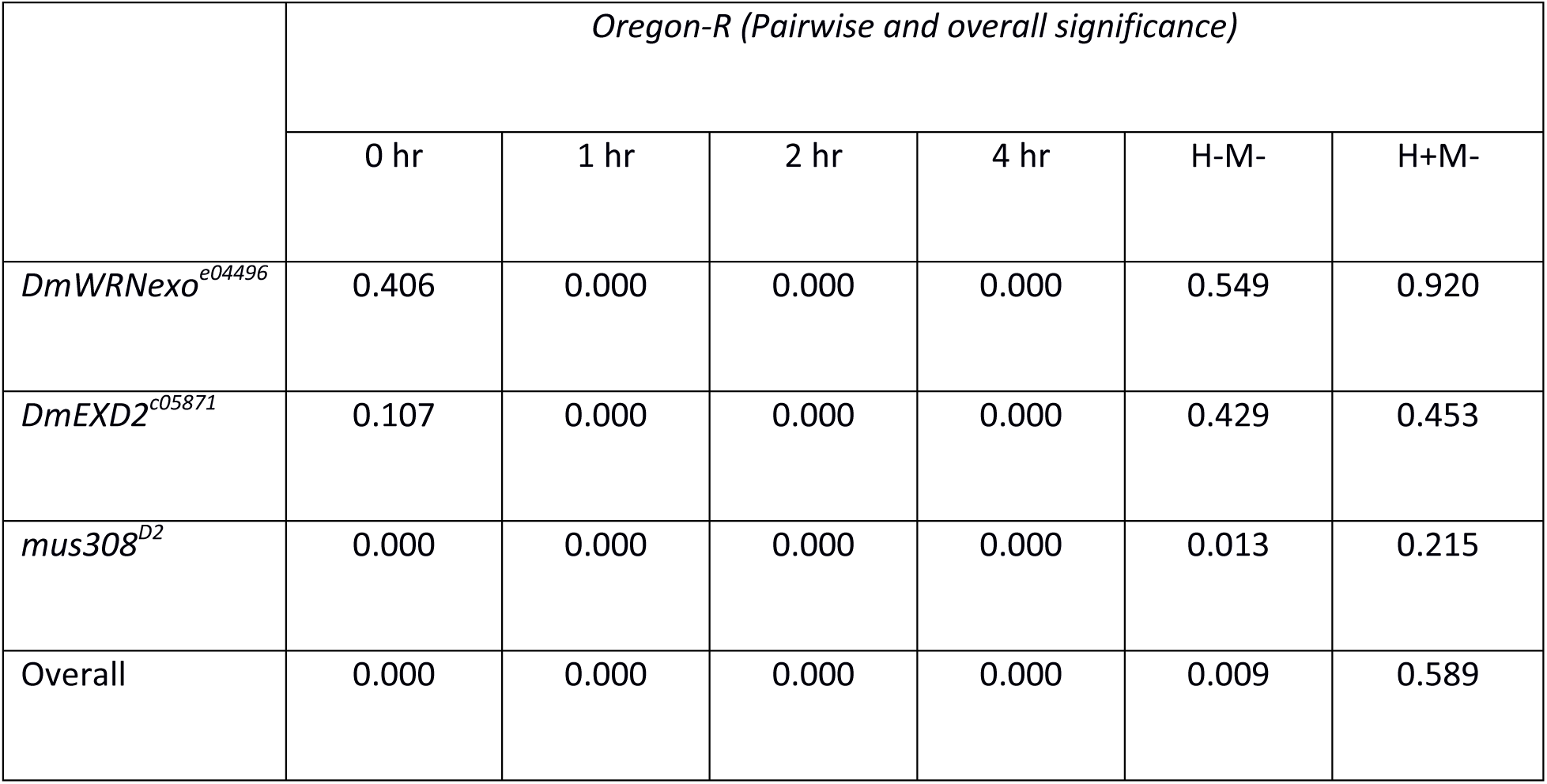
Statistical significance of repair efficiency of MMC-induced ICLs. Analysis of difference in comet tail moment at various recovery times following MMC-induced ICLs in fly larval neuroblasts in homozygous mutants versus using Oregon-R wild type controls. The Mann-Whitney U Wilcoxon W Test was used for pairwise comparisons between each mutant genotype and the wild type Oregon-R and overall significance tested using the Kruskal Wallis Test package, using SPSS. H-M- = control without either hydrogen peroxide (nicking) or MMC (cross-linking) treatment. H+M- = control with peroxide but without MMC. *** in Figure 6 indicates p<0.001.

